# WAT3R: Recovery of T-Cell Receptor Variable Regions From 3’ Single-Cell RNA-Sequencing

**DOI:** 10.1101/2022.01.26.477886

**Authors:** Marina Ainciburu, Duncan M. Morgan, Erica A. K. DePasquale, J. Christopher Love, Felipe Prósper, Peter van Galen

## Abstract

**Summary:** Diversity of the T-cell receptor (TCR) repertoire is central to adaptive immunity. The TCR is composed of α and β chains, encoded by the *TRA* and *TRB* genes, of which the variable regions determine antigen specificity. To generate novel biological insights into the complex functioning of immune cells, combined capture of variable regions and single-cell transcriptomes provides a compelling approach. Recent developments enable the enrichment of *TRA* and *TRB* variable regions from widely used technologies for 3’-biased single-cell RNA-sequencing (scRNA-seq). However, a comprehensive computational pipeline to process TCR-enriched data from 3’ scRNA-seq is not available. Here we present an analysis pipeline to process TCR variable regions enriched from 3’ scRNA-seq cDNA. The tool reports *TRA* and *TRB* nucleotide and amino acid sequences linked to cell barcodes, enabling the reconstruction of T-cell clonotypes with associated transcriptomes. We demonstrate the software using peripheral blood mononuclear cells (PBMCs) from a healthy donor and detect TCR sequences in a high proportion of single T-cells. Detection of TCR sequences is negligible in non-T-cell populations, demonstrating specificity. Finally, we show that TCR clones are larger in CD8 Memory T-cells than other T-cell types, indicating an association between T-cell clonotypes and differentiation states.

**Availability and implementation:** The Workflow for Association of T-cell receptors from 3’ single-cell RNA-seq (WAT3R), including test data, is available on GitHub (https://github.com/mainciburu/WAT3R), Docker Hub (https://hub.docker.com/r/mainciburu/wat3r), and a workflow on the Terra platform (https://app.terra.bio). The test dataset is available on GEO (accession number pending).

## 1. Introduction

During T-cell development, a series of recombinations shape the α and β chains that comprise the TCR, giving rise to 10^15^-10^21^ potential TCRs (La Gruta *et al*., 2018). The recombinations occur in the variable regions of the *TRA* and *TRB* genes that encode the TCR α/β chains. *TRA* and *TRB* determine T-cell specificity by shaping TCR recognition of antigens presented by MHC molecules. The recombined sequences can also be used to track T-cell clonotypes. With the advent of single-cell sequencing technologies, simultaneous capture of *TRA* and *TRB* sequences combined with transcriptional states provides a powerful approach to study T-cell biology (Stubbington *et al*., 2016).

Several protocols have been developed to combine scRNA-seq with recovery of TCR sequences, using cell barcodes to integrate both layers of information. Low-throughput single-cell methods performed on multiwell plates allow for the recovery of the complete *TRA* and *TRB* genes (Sade-Feldman *et al*., 2018; Stubbington *et al*., 2016). On the other hand, high-throughput methods based on microfluidics or microwell chips create libraries that can be enriched for TCR sequences (Oliveira *et al*., 2021; Zemmour *et al*., 2018; Oh *et al*., 2020). Among the latter, T-cell Receptor Enrichment to linK clonotypes (TREK-seq) is a recently published protocol to perform both transcript and TCR sequencing from 10x Genomics 3’ Gene Expression libraries (Miller *et al*., 2022; Tu *et al*., 2019). Although analysis tools exist for TCR bulk analysis (Bolotin *et al*., 2015) and 5’ single-cell protocols (CellRanger), there is a need for bioinformatics tools that facilitate the analysis of TCR variable regions enriched from 3’ scRNA-seq cDNA. Here, we describe WAT3R (pronounced “water”), an integrated pipeline that covers TCR-enriched data from preprocessing FASTQ files to alignment and the identification of T-cell clones.

## 2. Description

In accordance with the TREK-seq protocol, sequencing data is provided as two compressed FASTQ files. One contains the cell barcode and unique molecular identifier (UMI) sequences and the other the TCR sequence. First, files are reformatted to join the barcode, UMI, and corresponding TCR sequence in a single FASTQ. Next, the user can specify whether barcode and UMI correction should be performed. Barcode correction allows for one mismatch with barcodes in the 10x Single Cell 3’ v3 list (or a custom list provided by the user). UMI correction is performed by clustering together UMIs with one mismatch and considering the most abundant UMI as the correct one. Every barcode and UMI sequence is then added to the corresponding FASTQ read header, to be used as an identifier. Next, we apply a quality filter to remove every read with an average quality score lower than indicated (default is qscore < 25, Figure 1, top left). To account for barcode swapping (i.e., incorrect barcode assigning), TCR sequences with an identical barcode and UMI are subjected to clustering based on sequence similarity, using the USEARCH algorithm (Edgar, 2010). The identity threshold to measure similarity can be set by the user (default is 0.9). To ensure high data consistency, only TCR sequences are kept if the most abundant cluster represents a large proportion of the reads (default is 0.5) and is substantially larger than the second most abundant cluster (default ratio is 2.0) (Figure 1, top right).

**Figure 1.**
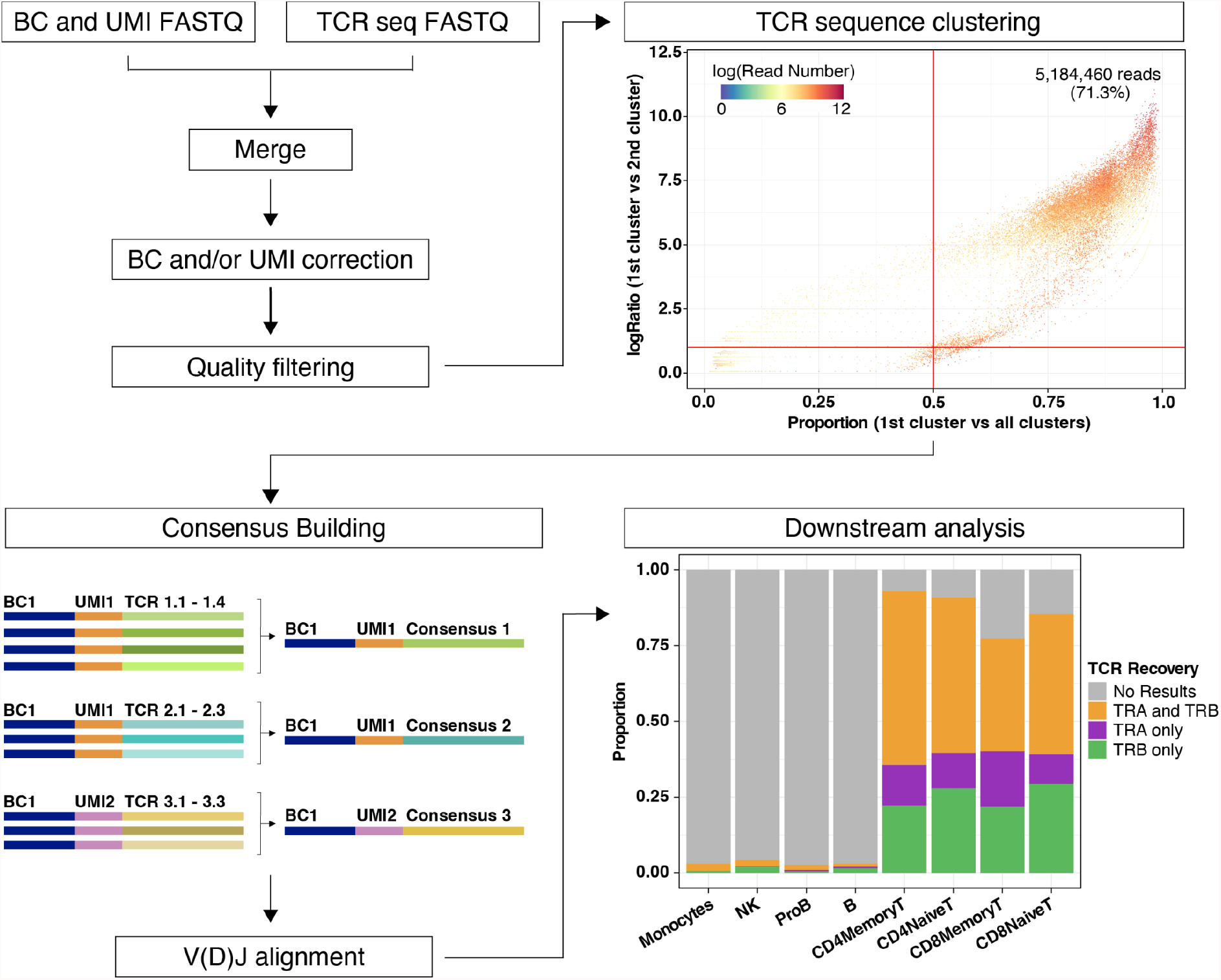
Overview of WAT3R. The workflow starts by merging two FASTQ files, correction of cell barcodes and UMIs, and quality filtering (top left). Clustering of TCR sequences with identical barcode and UMI is then performed. Top right dotplot shows evaluation of cluster quality by comparing the proportion of reads supporting the most abundant cluster (x-axis), the ratio of the most abundant cluster to the second (y-axis), and the number of reads supporting each TCR sequence with a specific barcode and UMI (color). Bottom left: TCR consensus sequences are generated and used for V(D)J alignment. In downstream analysis, results are integrated with a paired scRNA-seq dataset. Bottom right barplot shows the proportion of cells in the dataset, separated by cell type, for which WAT3R returned information on the *TRA* gene, *TRB* gene, or both.

Next, a consensus sequence is built for each of the clusters (Heiden *et al*., 2014). For a consensus to be constructed, we require a minimum of 3 reads and allow for a maximum error rate of 0.5 and a gap frequency of 0.5 per position; these parameters can be changed by the user (Figure 1, bottom left). Consensus sequences are aligned to the V(D)J segments reference provided by IMGT, using IgBLAST with the default parameters (Ye *et al*., 2013; Lefranc *et al*., 2015). This task is performed through the interface implemented in the Python package Change-O (Gupta *et al*., 2015). After selecting *TRA* and *TRB* sequences with the highest UMI counts, the CDR3 nucleotide/amino acid sequences and V(D)J calls are assigned to cell barcodes and saved in a results table.

As an optional step, the user can provide a file with cell barcodes and annotations coming from a paired scRNA-seq experiment to integrate with the *TRA* and *TRB* calls (Figure 1, bottom right). Overall, this pipeline returns two tables of results, one at the transcript level and another at the cell level. In addition, multiple quality control (QC) graphs and metrics are generated. WAT3R, together with the required software, reference data and documentation is available as a docker image. It is also available as a workflow on Terra which provides access to Google Cloud computing resources through a simple web-based user interface.

## 3. Results

We analyzed a human peripheral blood sample using 10x Genomics 3’ v3 scRNA-seq and TREK-seq to enrich *TRA* and *TRB* variable regions (Miller *et al*., 2022). The TREK-seq library was sequenced on a MiSeq to a depth of 8 million reads. After recovering 4.3% of the cell barcodes using the barcode correction algorithm, 98.3% of reads contained valid barcodes (Supplementary Figure 1). Likewise, 4.9% of the UMI sequences were corrected. Reads were filtered for an average q score above 25, which retained 92.3% of the original reads. For consensus building and subsequent alignment, 70.4% of the reads were valid. After removal of TCR sequence clusters below the proportion and ratio thresholds, 65.8% of the original reads were retained. These reads were used to generate a results table (Supplementary Figure 2).

We integrated these results with the paired 3’ scRNA-seq dataset with cell type annotations based on canonical marker genes (Supplementary Figure 3) (Hao *et al*., 2021). Detection of *TRA* or *TRB* sequences in 90% of all T-cells demonstrates efficient enrichment from the cDNA. In contrast, *TRA* or *TRB* sequences are present in only 3% of non-T-cells, demonstrating specificity. The *TRB* variable region is most efficiently enriched: *TRA* sequences are detected in 65% of single T-cells, *TRB* in 77%, and *TRA+TRB* in 52% (Figure 1, bottom right). As expected in a healthy individual, we did not observe any expanded T-cell clones dominating the sample. Nonetheless, the largest detected clones belong mainly to CD8 Memory T-cell subsets, in accordance with previous findings (Supplementary Figure 4) (DePasquale *et al*., 2021; Penter *et al*., 2021).

## 4. Conclusions

WAT3R is a comprehensive pipeline for the analysis of *TRA* and *TRB* variable regions enriched from the cDNA of widely used 3’ scRNA-seq protocols. From sequencing error correction, alignment, and quality controls to intersection with cell type annotations from scRNA-seq, this tool applies state-of-the-art algorithms to reliably detect T-cell clonotypes and initiate new discoveries in immunology.

## Acknowledgments

We thank the healthy donor for donating peripheral blood cells. We thank Julia Verga, Tyler Miller, Martin Villanueva, Charles Couturier, Daniel Ssozi, Jonathan Good, Jenny Noel and Alex Shalek for help with the TREK-seq protocol development and Yoke Seng Lee and Antonia Kreso for helpful feedback.

## Supplementary Data

**Supplementary Figure 1.**
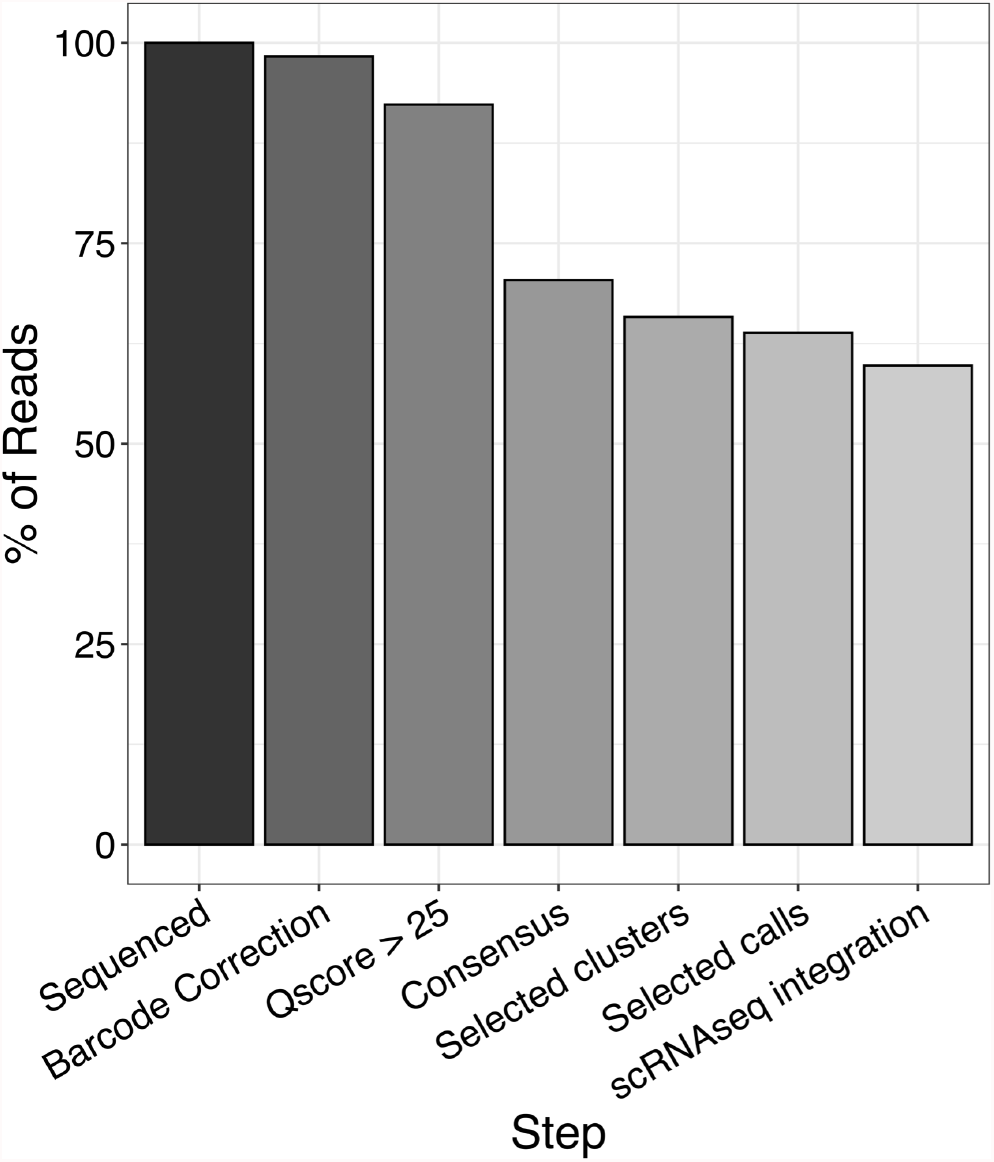
Retainment of raw sequencing reads. Barplot shows the percentage of reads passing the different filtering steps throughout the pipeline.

**Supplementary Figure 2.**
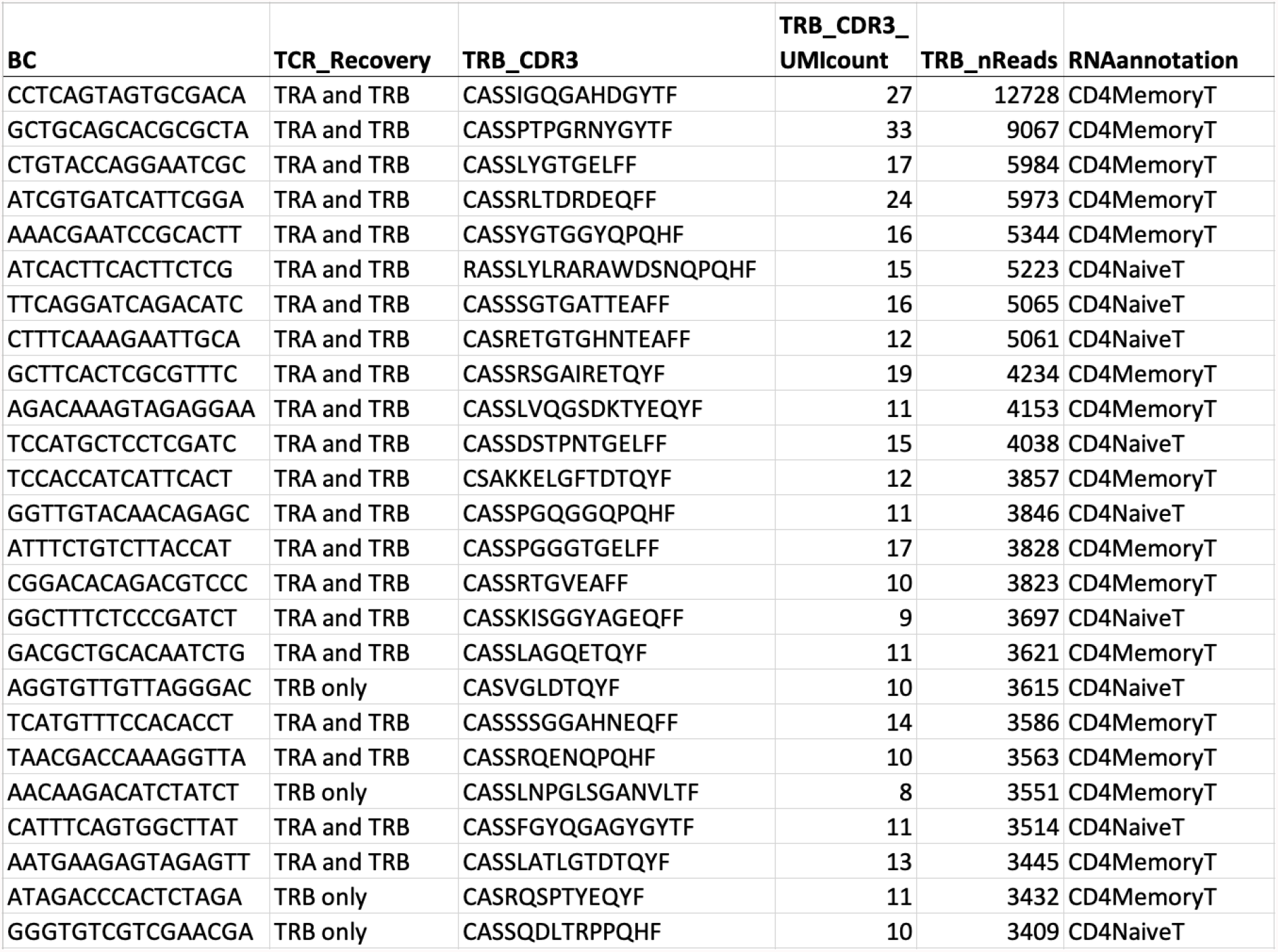
Excerpt of the results table. The complete results table contains additional columns and rows, including CDR3 loop nucleotide sequences, predicted V(D)J segments and *TRAV* calls. BC: cell barcode, TCR_Recovery: *TRA, TRB* or both, TRB_CDR3: predicted amino acid sequence of the CDR3 loop, TRB_CDR3_UMIcount: number of UMIs (transcripts) supporting the *TRB* call, TRB_nReads: number of reads supporting the *TRB* call, RNAannotation: cell type annotation from scRNA-seq.

**Supplementary Figure 3.**
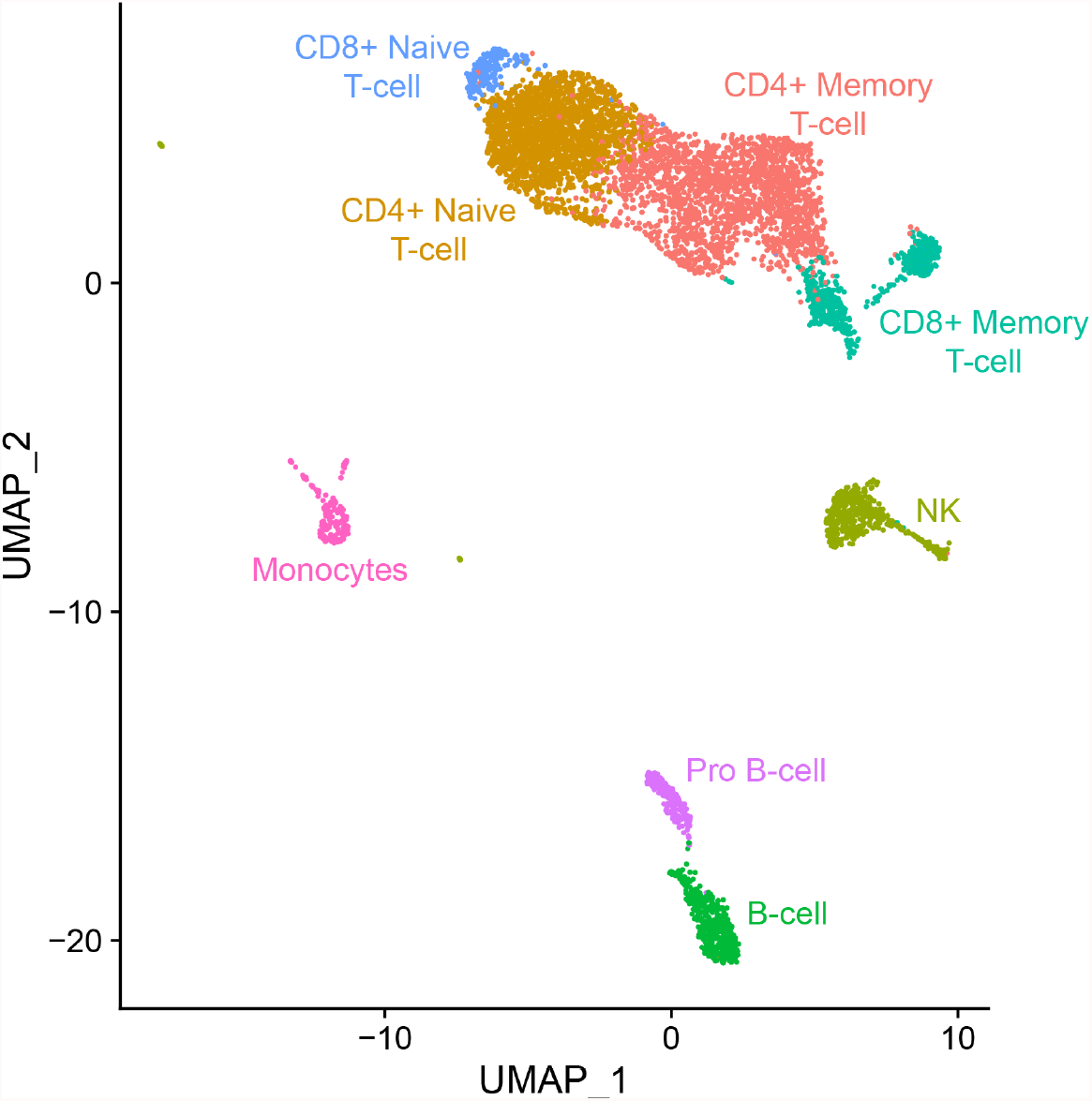
UMAP with cell type annotations. UMAP shows 6,300 cells from a healthy PBMC donor along with cell type annotations. Dimensionality reduction and clustering was performed using standard procedures (Hao *et al*., 2021). The cell type annotations correspond to the bottom right panel of Figure 1.

**Supplementary Figure 4.**
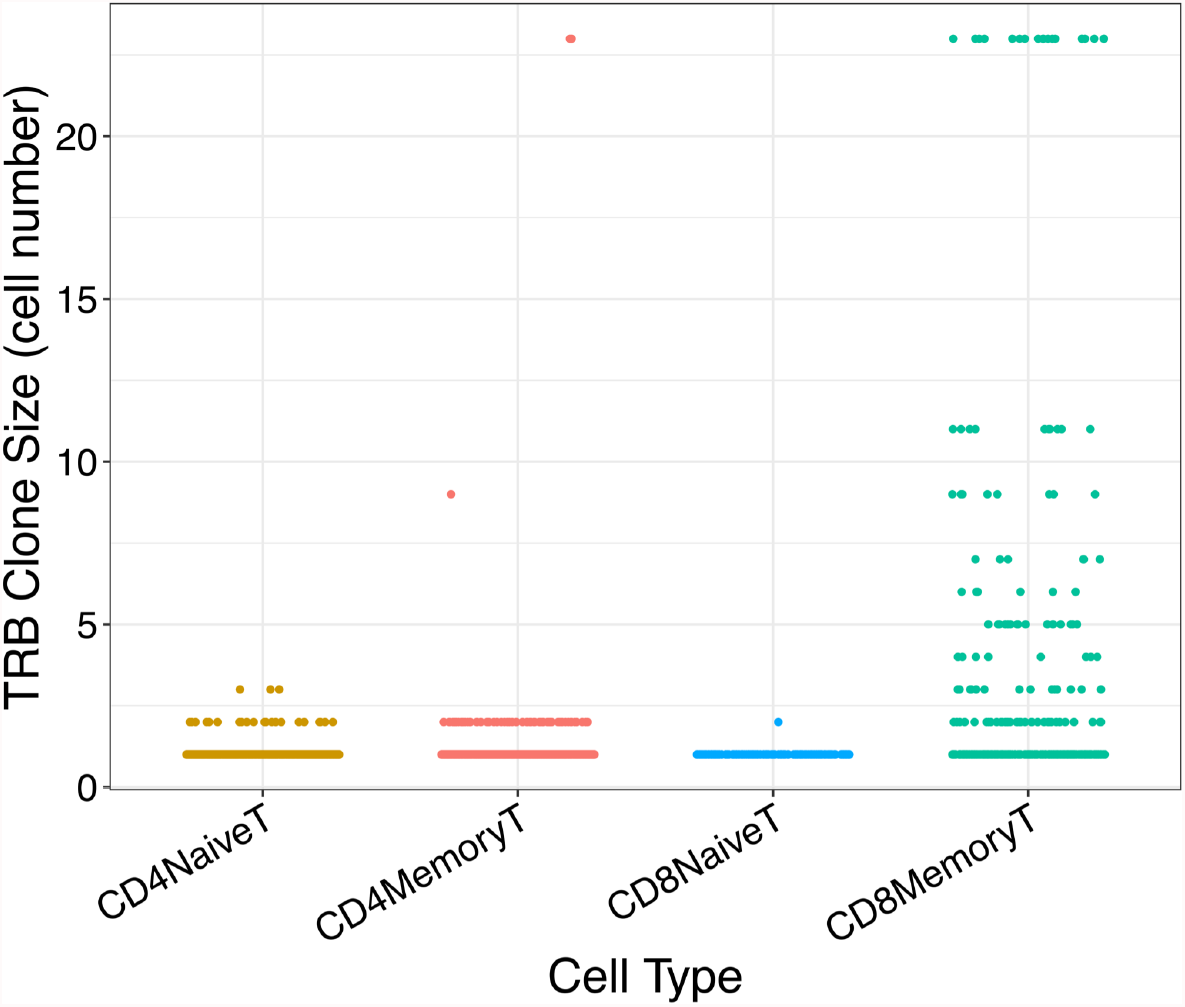
T-cell clonotype sizes are largest in CD8 Memory T-cells. Dot plot shows T-cell clone sizes (number of cells with the same *TRB* sequence) in four different T-cell populations from the healthy PBMC donor. Clone sizes are significantly larger in CD8 Memory T-cells compared to any of the other populations (*P* < 0.05).

## References

Bolotin, D.A. et al. (2015) MiXCR: software for comprehensive adaptive immunity profiling. Nat. Methods, 12, 380–381.

DePasquale, E.A.K. et al. (2021) Single-cell Multiomics Reveals Clonal T-cell Expansions and Exhaustion in Blastic Plasmacytoid Dendritic Cell Neoplasm. bioRxiv, 2021.12.01.470599.

Edgar, R.C. (2010) Search and clustering orders of magnitude faster than BLAST. Bioinformatics, 26, 2460–2461.

Gupta, N.T. et al. (2015) Change-O: a toolkit for analyzing large-scale B cell immunoglobulin repertoire sequencing data. Bioinformatics, 31, 3356–3358.

Hao, Y. et al. (2021) Integrated analysis of multimodal single-cell data. Cell, 184, 3573–3587.e29.

Heiden, J.A.V. et al. (2014) pRESTO: a toolkit for processing high-throughput sequencing raw reads of lymphocyte receptor repertoires. Bioinformatics, 30, 1930–1932.

La Gruta, N.L. et al. (2018) Understanding the drivers of MHC restriction of T cell receptors. Nat. Rev. Immunol., 18, 467–478.

Lefranc, M.-P. et al. (2015) IMGT®, the international ImMunoGeneTics information system® 25 years on. Nucleic Acids Res., 43, D413–22.

Miller, T.E. et al. (2022) Mitochondrial variant enrichment from high-throughput single-cell RNA-seq resolves clonal populations. Nat. Biotechnol.

Oh, D.Y. et al. (2020) Intratumoral CD4+ T Cells Mediate Anti-tumor Cytotoxicity in Human Bladder Cancer. Cell, 181, 1612–1625.e13.

Oliveira, G. et al. (2021) Phenotype, specificity and avidity of antitumour CD8+ T cells in melanoma. Nature, 596, 119–125.

Penter, L. et al. (2021) Coevolving JAK2V617F+ relapsed AML and donor T cells with PD-1 blockade after stem cell transplantation: an index case. Blood Adv.

Sade-Feldman, M. et al. (2018) Defining T Cell States Associated with Response to Checkpoint Immunotherapy in Melanoma. Cell, 175, 998–1013.e20.

Stubbington, M.J.T. et al. (2016) T cell fate and clonality inference from single-cell transcriptomes. Nat. Methods, 13, 329–332.

Tu, A.A. et al. (2019) TCR sequencing paired with massively parallel 3′ RNA-seq reveals clonotypic T cell signatures. Nat. Immunol., 20, 1692–1699.

Ye, J. et al. (2013) IgBLAST: an immunoglobulin variable domain sequence analysis tool. Nucleic Acids Res., 41, W34–40.

Zemmour, D. et al. (2018) Single-cell gene expression reveals a landscape of regulatory T cell phenotypes shaped by the TCR. Nat. Immunol., 19, 291–301.

